# Impact of Aging, Sex, and Species on the mRNA Expression of Matrix Metalloproteinases Following Ischemic Stroke

**DOI:** 10.64898/2026.02.24.707225

**Authors:** Siva Reddy Challa, Isidra M. Baker, Vishesh Vinayagam, Samantha N. Jackson, Nabeeha Khan, Sahil Reddy Mada, Pavani Unnam, Casimir A. Fornal, Jeffrey D. Klopfenstein, Krishna Kumar Veeravalli

**Affiliations:** Department of Cancer Biology and Pharmacology, University of Illinois College of Medicine at Peoria, Peoria, Illinois, USA; The University of Texas at Dallas, Richardson, Texas, USA; University of Maryland Baltimore County, Baltimore, Maryland, USA; Department of Neurosurgery, University of Illinois College of Medicine at Peoria, Peoria, Illinois, USA; Department of Pediatrics, University of Illinois College of Medicine at Peoria, Peoria, Illinois, USA; Department of Neurology, University of Illinois College of Medicine at Peoria, Peoria, Illinois, USA

**Keywords:** ischemia, reperfusion, stroke, matrix metalloproteinase, expression, species, age, sex

## Abstract

Matrix metalloproteinase (MMP) expression and function are highly context dependent, varying across physiological and pathological conditions. We previously documented the expression profiles of select MMPs in the ischemic brains of young male rodents. However, aging is a major risk factor for stroke in humans and is associated with vasculature alterations, increased oxidative stress, and elevated inflammation. In addition, sex differences have been reported in stroke incidence and severity. Despite this, the effects of age, sex, and species on brain MMP gene expression after cerebral ischemia/reperfusion (I/R) has not been systematically examined. Therefore, we investigated how age, sex, and species influence the mRNA expression of all known MMPs (22 total) in the brain following cerebral I/R. Moderate-to-severe neurological deficits were induced by transient middle cerebral artery occlusion (MCAO) followed by reperfusion in young and aged male and female C57BL/6 mice and in young male Sprague-Dawley rats. Brain tissue from the ipsilateral (ischemic) hemisphere was collected on post-MCAO day 1, and MMP mRNA levels were quantified by real-time PCR and expressed as fold change relative to the sham control group. Across species, MMP-3, MMP-8, MMP-12, MMP-13, MMP-19, MMP-20, and MMP-27 were upregulated in both rats and mice. Species-specific increases were also observed: MMP-1, MMP-7, MMP-9, MMP-14, MMP-21, and MMP-25 were upregulated only in rats, whereas MMP-10 was upregulated only in mice. The most strongly upregulated MMPs were MMP-12 in rats and MMP-3, MMP-10, and MMP-12 in mice. By contrast, MMP-15 and MMP-17 were downregulated in both species, whereas MMP-23 and MMP-24 were downregulated only in rats and mice, respectively. Within mice, MMP-3, MMP-10, MMP-12, MMP-19, MMP-20, and MMP-21 increased in both sexes and age groups, except for MMP-19 in aged males and MMP-21 in young males. MMP-14 increased only in females (young and aged), whereas MMP-27 increased only in males (young and aged). Notably, MMP-3, MMP-10, and MMP-12 were the three most highly upregulated MMPs in both male and female mice regardless of age. Overall MMP mRNA expression levels were higher in aged male mice and lower in aged female mice relative to sex-matched young mice. Among all MMPs examined, MMP-12 showed the most marked upregulation across species and, within mice, across age groups and sexes. Collectively, these findings demonstrate that brain MMP gene expression after cerebral I/R is modulated by age, sex, and species, underscoring the importance of incorporating these biological variables when targeting MMPs individually or in combination in preclinical rodent stroke models.

## 1. Introduction

Stroke is one of the leading causes of death and disability worldwide (Feigin et al., 2025). Ischemic stroke, resulting from the occlusion of cerebral blood vessels, accounts for approximately 87% of all stroke cases (Virani et al., 2020). Recanalization therapies, including thrombolysis with tissue-type plasminogen activator or tenecteplase and endovascular thrombectomy, restore cerebral blood flow and initiate reperfusion of the previously ischemic tissue. Although reperfusion is essential for salvaging ischemic tissue, it can also lead to secondary brain damage (referred to as reperfusion injury) through oxidative stress and mitochondrial dysfunction, excitotoxicity, cell death pathways, cerebral edema, blood-brain barrier (BBB) disruption, hemorrhagic transformation, and inflammation (Cheng et al., 2024). Thus, therapies that limit progressive injury and improve functional recovery after cerebral ischemia/reperfusion (I/R) are urgently needed.

Increased expression and/or activity of several matrix metalloproteinases (MMPs) after cerebral I/R is implicated in cerebral edema, BBB disruption, hemorrhage, leukocyte infiltration, neuroinflammation, apoptosis, increased infarct volume, and sensorimotor and cognitive deficits (Lee et al., 2004; Si-Tayeb et al., 2006; Zhao et al., 2006; Rosenberg and Yang, 2007; Wasserman and Schlichter, 2007; Cuadrado et al., 2009; Yang et al., 2010; Chelluboina and Klopfenstein et al., 2015; Chelluboina and Warhekar et al., 2015; Arruri et al., 2022; Challa et al., 2022). Although 23 MMPs have been identified to date in humans, gene sequences are available for only 22 MMPs in rodents, and not for MMP-26. While the expression of all known MMPs has not been comprehensively examined in ischemic stroke, increased levels of MMP-1, MMP-2, MMP-3, MMP-8, MMP-9, MMP-10, and MMP-13 have been reported in ischemic human brain tissue (Cuadrado et al., 2009). Previously, we reported the mRNA expression changes in numerous MMPs in the ischemic brains of young male rats and young male mice on post-MCAO day 1 after cerebral I/R (Chelluboina and Warhekar et al., 2015; Nalamolu et al., 2018). In animal models of ischemic stroke, MMP-9 has been extensively studied and consistently shown to exert deleterious effects; whereas potentially harmful or beneficial effects have been reported for MMP-3, MMP-8, MMP-10, MMP-12, and MMP-13 (Orbe et al., 2011; Chelluboina and Klopfenstein et al., 2015; Chelluboina and Warhekar et al., 2015; Hafez et al., 2016; Han et al., 2016; Ma et al., 2016; Roncal et al., 2017; Hirono et al., 2018; Arruri et al., 2022; Challa et al., 2022; Hamblin et al., 2024). Notably, these investigations were conducted predominantly in males and in young rodents, which are less clinically representative of the older populations most susceptible to ischemic stroke and poorer outcomes.

Aging is associated with structural and functional changes in cerebral vasculature, along with increased oxidative stress and inflammation (Sierra et al., 2011). Thus, age is a major non-modifiable risk factor for stroke. Stroke risk doubles with each decade after age 55, and approximately 75% of strokes occur in individuals over 65 (Mozaffarian et al., 2015). Additionally, sex-based distinctions in the incidence, severity, and outcomes of acute ischemic stroke (AIS) have been reported (Zou et al., 2017). Age significantly influences the manifestation of sexual dimorphism in ischemic stroke. For example, in both humans and animal models, younger females generally experience less brain damage and better outcomes, whereas older females demonstrate substantially greater damage and worse functional outcomes compared to age-matched males (Alkayed et al., 1998; Vannucci et al., 2001; Roy-O’Reilly and McCullough, 2018). However, the impact of aging and sex on the expression of all known MMPs in the ischemic brain following cerebral I/R remains unknown. Therefore, the aim of this study was not only to assess the impact of age and sex on the mRNA expression profiles of all known MMPs in the ischemic brain but also to evaluate the reproducibility of previously reported mRNA expression patterns of numerous MMPs in young male rats and mice. Characterizing MMP expression across age, sex, and species may help to identify biologically relevant targets, guide the development of effective therapeutic strategies, and support preclinical testing in clinically relevant stroke models to improve translational potential.

## 2. Materials and Methods

### 2.1. *Animals*, procedures, and compliance

C57BL/6J mice were obtained from The Jackson Laboratory (Bar Harbor, ME, USA), and Sprague-Dawley (SD) rats were obtained from Envigo (Indianapolis, IN, USA). Animals were housed in the University of Illinois College of Medicine Peoria (UICOMP) Laboratory Animal Care Facility under controlled temperature and humidity on a 12-h light-dark cycle, with *ad libitum* access to food and water. A total of 60 C57BL/6J mice (both sexes combined; young [3-4 months old] and aged [17-19 months old]) and 14 young (3-month-old) male SD rats were used in this study and were randomly assigned to either Control or Stroke groups (Supplementary Table 1). Cerebral I/R was induced by right transient middle cerebral artery occlusion (MCAO) using the intraluminal monofilament suture technique in mice (1-h MCAO) and rats (2-h MCAO), as described recently by our group (Challa et al., 2026). Pre- and post-surgical care was provided in accordance with the IMPROVE guidelines (Ischemia Models: Procedural Refinements of In Vivo Experiments) (Percie du Sert et al., 2017). Modified Neurological Severity Score (mNSS) assessment in rats and Neurological Deficit Score (NDS) assessment in mice were performed on post-MCAO day 1 to evaluate stroke severity, as described recently by our group (Challa et al., 2026). Rats with mNSS < 8 and mice with NDS scores < 2 on post-MCAO day 1 were excluded. In addition, animals that died or were euthanized during the study period, as well as those exhibiting post-mortem bleeding in the region of the MCA, were excluded from analysis. All animal procedures were conducted in accordance with a protocol approved (Protocol #1284641) by the Institutional Animal Care and Use Committee at UICOMP. All experiments adhered to the scientific, humane, and ethical principles of UICOMP and to the guidelines outlined in the *Guide for the Care and Use of Laboratory Animals* (NIH Publication No. 86-23, revised; U.S. Department of Health and Human Services). Animal experiments were designed, conducted, and reported in accordance with the Animal Research: Reporting of In Vivo Experiments (ARRIVE) guidelines (Percie du Sert et al., 2020).

### 2.2. Brain tissue collection, RNA isolation, and cDNA synthesis

On post-MCAO day 1, brain tissue was collected from young and aged male and female mice and from young male rats in the control and stroke groups. Under deep isoflurane anesthesia, animals were perfused intracardially with ice-cold 1× phosphate-buffered saline (PBS) and brains were rapidly removed, and the ipsilateral (ischemic) hemisphere was separated from the contralateral hemisphere and stored at -80 °C until RNA extraction. Total RNA was extracted from the entire ipsilateral cerebral hemisphere using TRIzol Reagent (Invitrogen, Carlsbad, CA, USA). One microgram of total RNA from each sample was reverse transcribed into cDNA using the iScript cDNA Synthesis Kit (Bio-Rad Laboratories, Hercules, CA, USA). The resulting cDNA was diluted 1:10 in nuclease free water and stored at -20 °C for subsequent analysis.

### 2.3. Real-time PCR analysis

For each cDNA sample, reactions were assembled using iTaq Universal SYBR Green Supermix (Bio-Rad Laboratories, Hercules, CA, USA). Forward and reverse primers (Supplementary Tables 2 and 3) for the target genes (Integrated DNA Technologies, Coralville, IA, USA) were diluted 1:10 in nuclease-free water. PCR reactions were performed in triplicates using the following thermal cycling conditions: initial denaturation at 95 °C for 5 min; 40 cycles of 95 °C for 30 sec, 59-62 °C for 30 sec, and 72 °C for 30 sec; and a final extension at 72 °C for 5 min. Reactions were run on an iCycler IQ Multi-Color Real-Time PCR Detection System (Bio-Rad Laboratories, Hercules, CA, USA). Data were collected using iCycler IQ software (Bio-Rad Laboratories, Hercules, CA, USA) and expressed as threshold cycle (Ct) values. For each sample, the mean Ct values of the triplicates, after removal of any outlier, was used for analysis. *18S* rRNA served as the internal reference gene. Relative target gene expression was normalized to *18S* rRNA, and fold change (test vs. control) was calculated as 2^(ΔCt control)/2^(ΔCt test), where ΔCt = Ct (target) - Ct (*18S* rRNA).

### 2.4. Statistical analysis

Statistical analyses were performed using GraphPad Prism 10.4.2 for Windows (GraphPad Software, San Diego, CA, USA). Outliers were identified using the ROUT test (Q = 1%) and excluded from analysis. Quantitative data were tested for normality (Shapiro-Wilk and Kolmogorov-Smirnov tests) and equality of variance (F-test). Based on the outcomes of these tests, appropriate statistical tests were applied, including the Mann-Whitney test when normality was not met, and for normally distributed data, a two-tailed unpaired t-test with Welch’s correction when variances were unequal. Differences between groups were considered significant at *p* < 0.05. All data are expressed as mean ± SEM.

## 3. Results

### 3.1. Post-stroke mortality, weight loss, and neurological deficits

A total of 2 rats and 12 mice were excluded from the study (Table 1). On post-MCAO day 1, stroke-induced rats included in the analysis had modified Neurological Severity Scores (mNSS) ranging from 8 to 12, consistent with a moderate injury (Supplementary Table 4). Similarly, mice had Neurological Deficit Scores (NDS) of 2-3, indicating a moderate-to-severe injury (Supplementary Table 5). On post-MCAO day 1, percent body weight loss relative to baseline was significantly greater in young male (*p* = 0.001) and young female mice (*p* = 0.0035) compared with their respective control groups (Supplementary Fig. 1A). In contrast, percent weight loss did not differ significantly between control and stroke groups in aged male or aged female mice (Supplementary Fig. 1B). Consistent with the findings in young mice, stroke-induced young rats exhibited significantly greater percent weight loss than controls (*p* = 0.0026) (Supplementary Fig. 2).

### 3.2. Overall baseline expression levels of each MMP within each age and sex group

In the four groups of control mice (young male, young female, aged male, and aged female), baseline expression of each MMP was evaluated within each group by comparing its normalized Ct value to the group mean normalized Ct calculated across all measured MMPs. The mean normalized Ct values (averaged across all MMPs) were 21.14 in young males, 19.69 in young females, 20.58 in aged males, and 19.71 in aged females (Supplementary Fig. 3). Within age-by-sex group, we identified MMPs with normalized Ct values ≥ 15% above or ≤ 15% below the group-specific mean. This threshold corresponds to an approximately 3-Ct deviation (∼8-fold difference in relative mRNA levels), with higher Ct indicating substantially lower expression and lower Ct indicating substantially higher expression. In young mice, MMP-8, MMP-12, and MMP-27 showed lower baseline expression in both sexes, whereas MMP-16, MMP-17, and MMP-24 showed higher baseline expression in both sexes; MMP-14 and MMP-15 exhibited higher baseline expression in males only (Supplementary Fig. 3A). Consistent with these patterns, aged mice also exhibited lower baseline expression of MMP-8, MMP-12, and MMP-27, and higher baseline expression of MMP-16, MMP-17, and MMP-24 in both sexes (Supplementary Fig. 3B).

### 3.3. Impact of aging on the expression of MMPs in the ischemic brains of mice

Ischemic stroke was associated with upregulation of multiple MMPs in the ipsilateral (ischemic) hemisphere. Aging influenced both the magnitude and pattern of this response in a sex-dependent manner. Overall, aged males generally exhibited higher expression levels of many MMPs, whereas aged females generally exhibited lower expression levels, relative to sex-matched young mice (Fig. 1).

**Figure 1.**
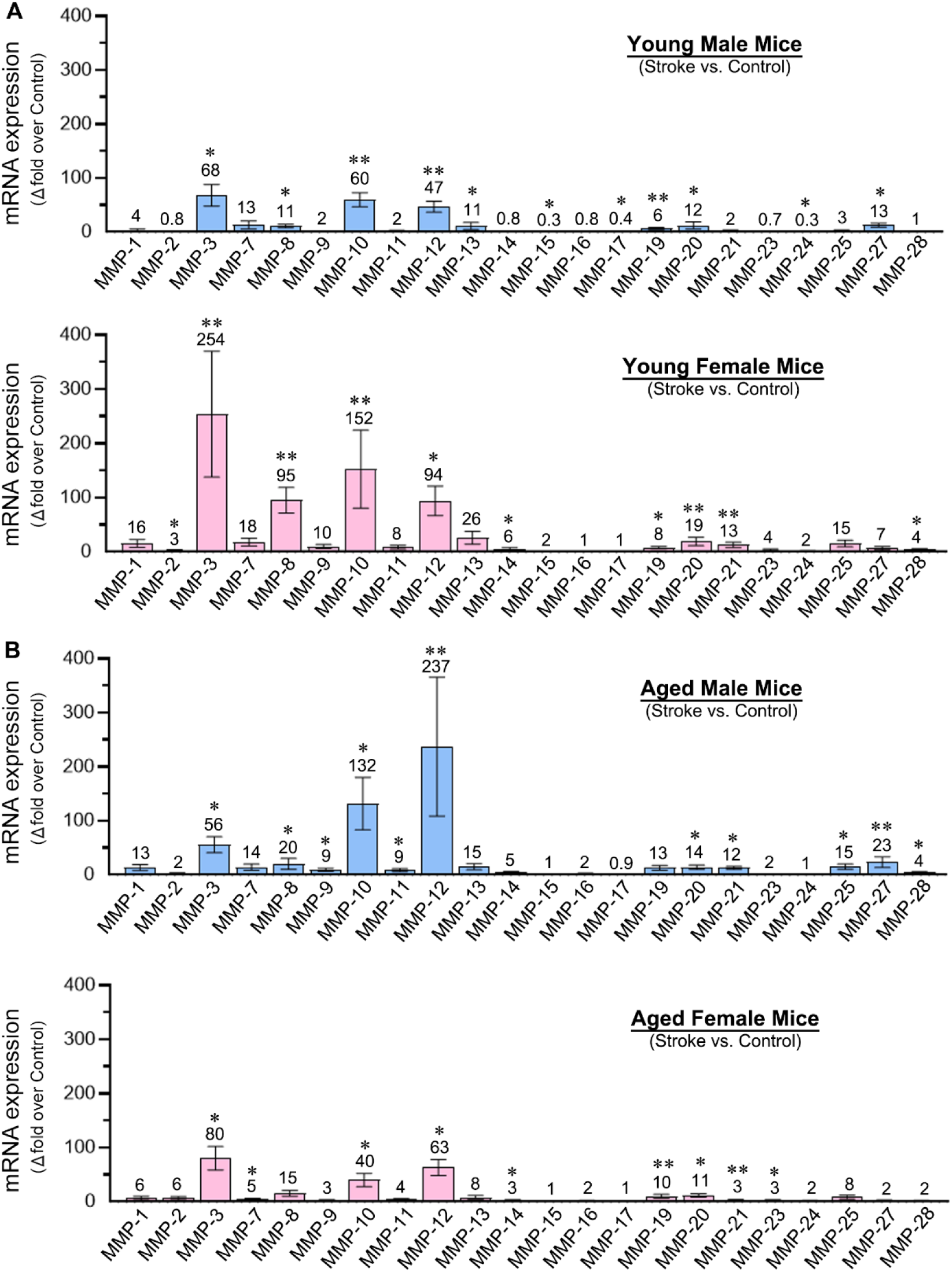
MMP expression in the mouse brain following ischemic stroke in young and aged mice of both sexes. Bar graphs show the quantified mRNA expression levels (fold change vs. the respective control group) for all mouse MMPs in the ipsilateral (ischemic) hemisphere of (**A**) young male and female mice and (**B**) aged male and female mice subjected to 1-h MCAO followed by reperfusion. Values above each bar indicate the fold change. Data are presented as mean ± SEM; n = 5–6/group. \**p* < 0.05, \*\**p* < 0.01 versus control group.

In males, stroke-induced changes in MMP expression were statistically significant for many MMPs in both age groups (Fig. 1A, B). Among the MMPs that were significantly upregulated, MMP-3, MMP-8, MMP-10, MMP-12, MMP-20, and MMP-27 increased in both young and aged stroke groups. By contrast, MMP-13 and MMP-19 were upregulated only in young mice, whereas MMP-9, MMP-11, MMP-21, MMP-25, and MMP-28 were upregulated only in aged mice (Fig. 2A). Compared with controls, increases of 40-fold or greater in MMP expression were observed for MMP-3 (68-fold; *p* = 0.0208), MMP-10 (60-fold; *p* = 0.0022), and MMP-12 (47-fold; *p* = 0.0022) in young male mice, and MMP-3 (56-fold; *p* = 0.0332), MMP-10 (132-fold; *p* = 0.0441), and MMP-12 (237-fold; *p* = 0.0022) in aged male mice (Fig. 1).

**Figure 2.**
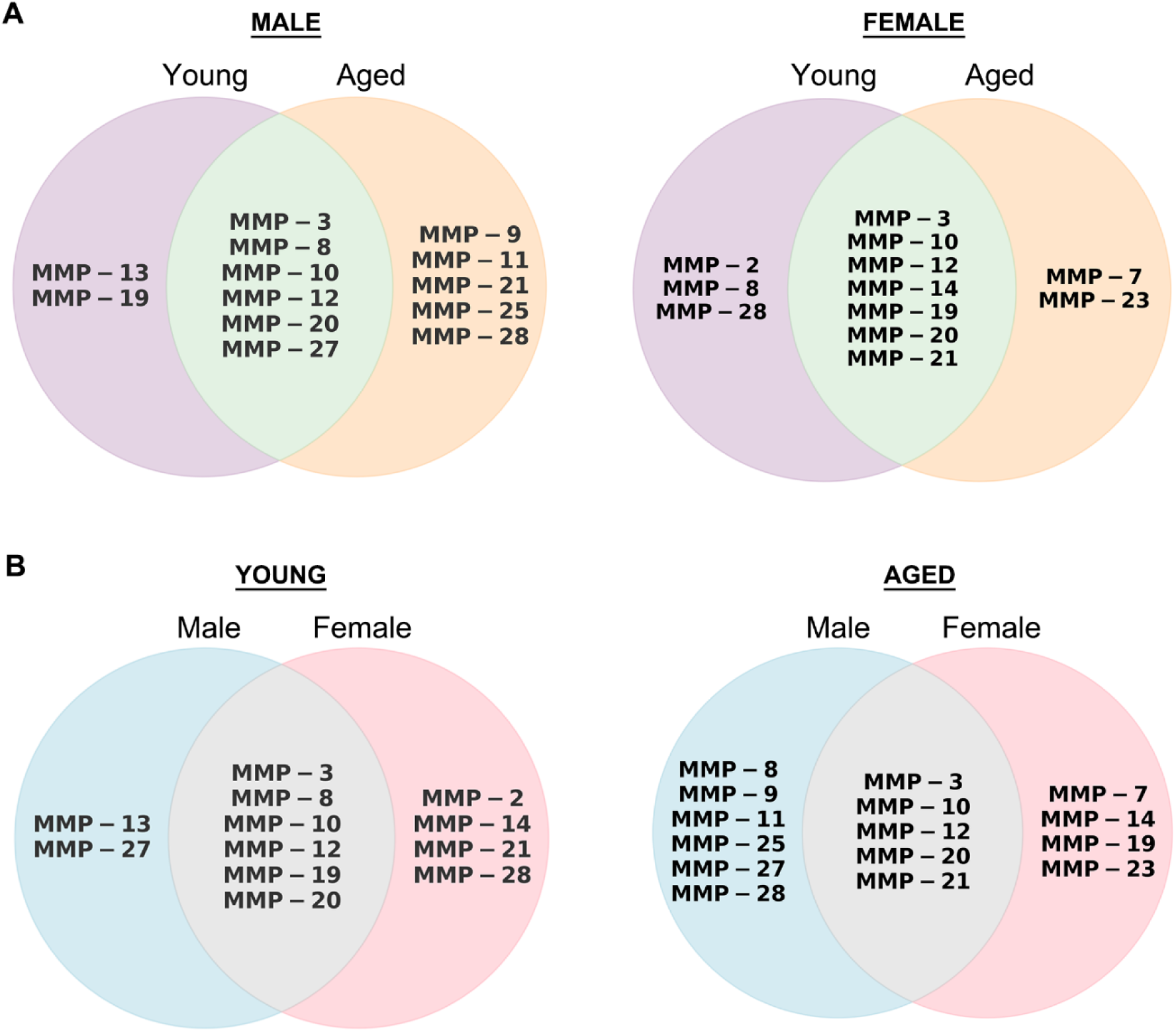
Similarities and distinct variations in MMP expression in the brain after ischemic stroke as a function of age and sex. Venn diagrams illustrate the similarities and differences among significantly upregulated MMPs (vs. controls) post-MCAO and reperfusion in the ipsilateral (ischemic) hemisphere of mice, grouped by (**A**) age and (**B**) sex, across the 22 MMPs examined.

In females, as in males, stroke-induced changes in MMP expression were statistically significant for many MMPs in both age groups (Fig. 1A, B). Among the MMPs that were significantly upregulated, Although MMP-3, MMP-10, MMP-12, MMP-14, MMP-19, MMP-20, and MMP-21 increased in both young and aged stroke groups. By contrast, MMP-2, MMP-8, and MMP-28 were upregulated only in young mice, whereas MMP-7 and MMP-23 were upregulated only in aged mice (Fig. 2A). Compared with controls, increases of 40-fold or greater in MMP expressions were observed for MMP-3 (254-fold; *p* = 0.0022), MMP-8 (95-fold; *p* = 0.0022), MMP-10 (152-fold; *p* = 0.0022), and MMP-12 (94-fold; *p* = 0.0192) in young female mice, and MMP-3 (80-fold; *p* = 0.0223), MMP-10 (40-fold; *p* = 0.0355), and MMP-12 (63-fold; *p* = 0.0137) in aged female mice (Fig. 1).

Given the observed age-related differences in post-stroke MMP expression profiles within the ischemic brain after cerebral I/R, we next examined the influence of age on baseline MMP expression in control mice. In males, aging significantly increased baseline expression of several MMPs, including MMP-1 (4.4-fold; *p* = 0.0315), MMP-7 (4-fold; *p* = 0.0196), MMP-8 (5.6-fold; *p* = 0.0173), MMP-12 (2.9-fold; *p* = 0.0168), MMP-13 (2.7-fold; *p* = 0.024), MMP-25 (8.7-fold; *p* = 0.0032), and MMP-27 (2.2-fold; *p* = 0.0124), whereas MMP-15 was reduced (0.5-fold; *p* = 0.0175) (Fig. 3A). In females, aging did not increase baseline MMP expression; instead, it was associated with decreased expression of MMP-2 (0.3-fold; *p* = 0.0003) and MMP-24 (0.7-fold; *p* = 0.0173) (Fig. 3B).

**Figure 3.**
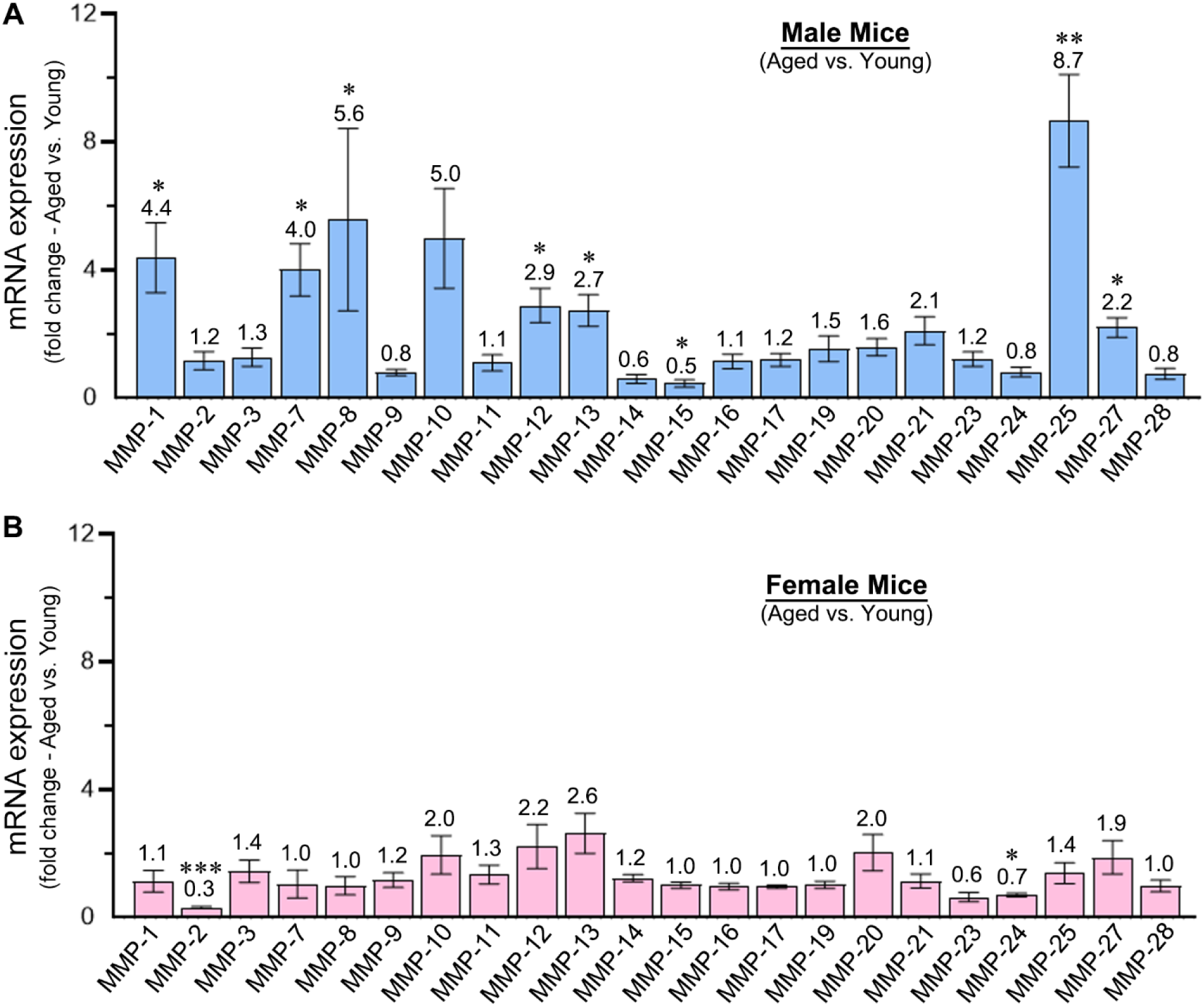
Impact of aging on MMP expression in the ischemic brain of male and female mice. Bar graphs show the quantified baseline mRNA expression levels as fold change (old vs. young) for various MMPs in the ipsilateral (ischemic) hemisphere of (**A**) male and (**B**) female mice subjected to 1-h MCAO followed by reperfusion. Values above each bar indicate the fold change. Data are presented as mean ± SEM; n = 5–6/group. \**p* < 0.05, \*\**p* < 0.01, \*\*\**p* < 0.001, old vs. young mice.

### 3.4. Sex differences and similarities in the expression of MMPs after cerebral I/R in mice

In young mice, females generally exhibited higher expression levels of numerous MMPs in the ipsilateral (ischemic) hemisphere compared with age-matched males (Fig. 1A). In contrast, in aged mice, females generally exhibited lower expression levels of many MMPs relative to age-matched males (Fig. 1B).

In young mice, MMP-3, MMP-8, MMP-10, MMP-12, MMP-19, and MMP-20 were upregulated in both sexes. By contrast, MMP-13 and MMP-27 were upregulated only in males, whereas MMP-2, MMP-14, MMP-21, and MMP-28 were upregulated only in females (Fig. 2B). In aged mice, MMP-3, MMP-10, MMP-12, MMP-20, and MMP-21 were upregulated in both sexes. By contrast, MMP-8, MMP-9, MMP-11, MMP-25, MMP-27, and MMP-28 were upregulated only in males, whereas MMP-7, MMP-14, MMP-19, and MMP-23 were upregulated only in females (Fig. 2B).

Considering the sex-dependent differences observed in post-stroke MMP mRNA expression profiles in the ischemic brain, we next examined the influence of sex on baseline MMP expression in control mice across both age groups. Baseline expression in females was quantified as fold change relative to age-matched males within each age group. Baseline MMP expression levels were substantially higher in females than in age-matched males (Fig. 4). In young mice, baseline expression in females was significantly higher than males for 15 of 22 MMPs, including MMP-1 (7.6-fold; *p* = 0.0028), MMP-2 (3.6-fold; *p* = 0.0001), MMP-3 (2.7-fold; *p* = 0.0017), MMP-7 (8.8-fold; *p* < 0.0001), MMP-8 (3.3-fold; *p* = 0.0043), MMP-11 (3.5-fold; *p* = 0.0017), MMP-12 (4-fold; *p* = 0.0385), MMP-16 (2.1-fold; *p* = 0.0245), MMP-17 (2.1-fold; *p* = 0.0143), MMP-19 (3.5-fold; *p* = 0.0043), MMP-20 (2.7-fold; *p* = 0.0098), MMP-23 (3-fold; *p* = 0.011), MMP-25 (8.1-fold; *p* = 0.0025), MMP-27 (3.3-fold; *p* = 0.0008), and MMP-28 (2.7-fold; *p* = 0.002) (Fig. 4A). In aged mice, baseline expression in females was significantly higher than males for 8 of 22 MMPs, including MMP-3 (2.8-fold; *p* = 0.0326), MMP-9 (4.2-fold; *p* = 0.0121), MMP-11 (4.6-fold; *p* = 0.0185), MMP-14 (2.1-fold; *p* = 0.0187), MMP-16 (2-fold; *p* = 0.0232), MMP-17 (1.8-fold; *p* = 0.0075), MMP-19 (2.7-fold; *p* = 0.0064), and MMP-28 (4-fold; *p* = 0.0126) (Fig. 4B). Notably, none of the 22 MMPs showed lower baseline expression in females relative to age-matched males.

**Figure 4.**
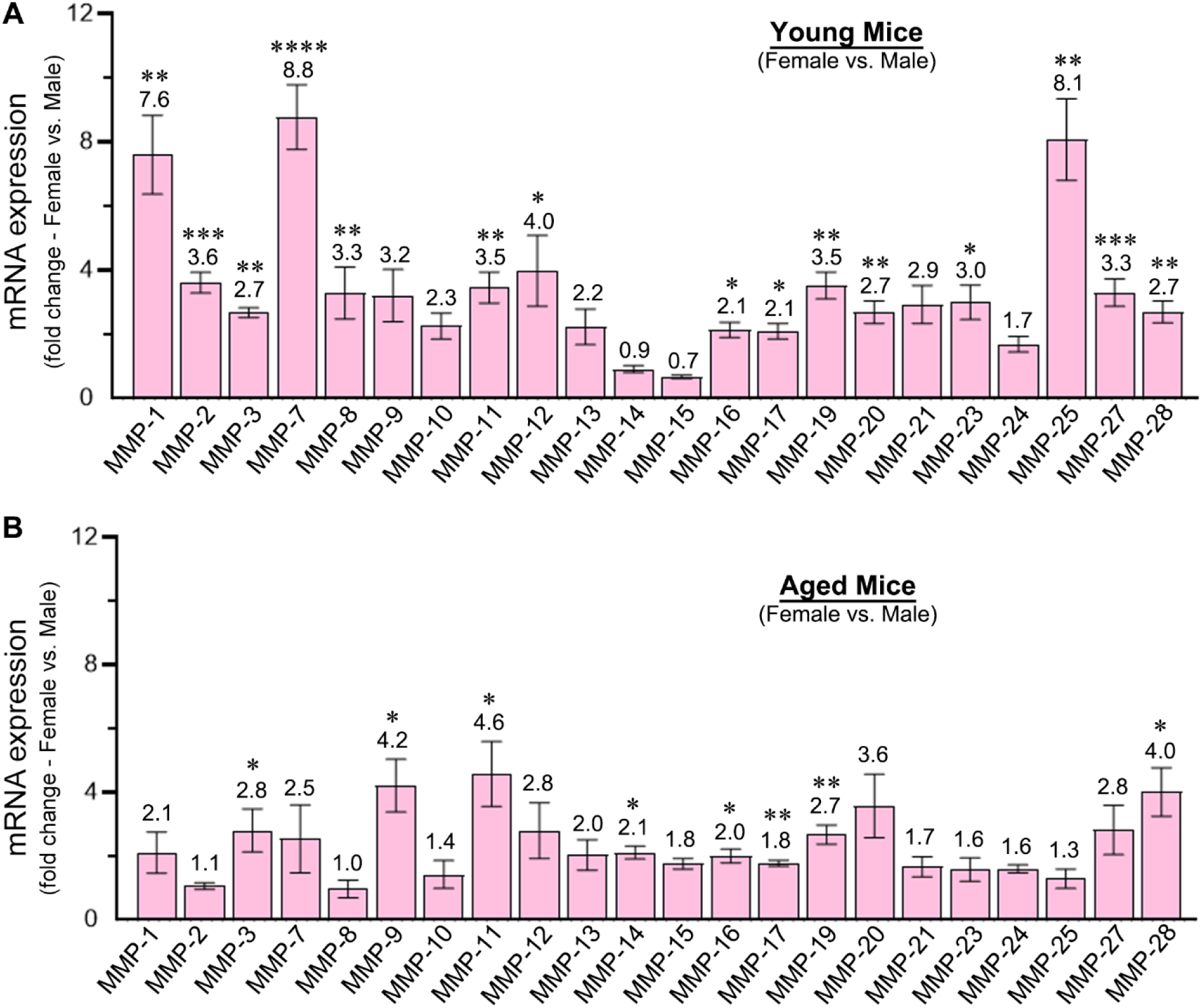
Impact of sex on MMP expression in the ischemic brain of young and aged mice. Bar graphs show the quantified baseline mRNA expression levels as fold change (female vs. age-matched male) for various MMPs in the ipsilateral (ischemic) hemisphere of (**A**) young and (**B**) aged mice subjected to 1-h MCAO followed by reperfusion. Values above each bar indicate the fold change. Data are presented as mean ± SEM; n = 5–6/group. \**p* < 0.05, \*\**p* < 0.01, \*\*\**p* < 0.001, \*\*\*\**p* < 0.0001, female vs. age-matched male.

### 3.5. Influence of species on the expression of MMPs following cerebral I/R

Cerebral I/R altered MMP mRNA expression in the ipsilateral (ischemic) hemisphere in both rats and mice (Fig. 5A). In both rats and mice, stroke-induced changes were statistically significant for most MMPs. Across species, MMP-3, MMP-8, MMP-12, MMP-13, MMP-19, MMP-20, and MMP-27 were upregulated in both rats and mice. Species-specific upregulation was also observed: MMP-1, MMP-7, MMP-9, MMP-14, MMP-21, and MMP-25 were upregulated only in rats, whereas MMP-10 was upregulated only in mice (Fig. 5B). The most strongly upregulated MMP in rats was MMP-12 (46-fold; *p* = 0.001). By contrast, the most strongly upregulated MMPs in mice were MMP-3 (68-fold; *p* = 0.0208), MMP-10 (60-fold; *p* = 0.0022), and MMP-12 (47-fold; *p* = 0.0022). Finally, MMP-15 and MMP-17 were downregulated in both species, whereas MMP-23 and MMP-24 were downregulated only in rats and mice, respectively (Fig. 5C).

**Figure 5.**
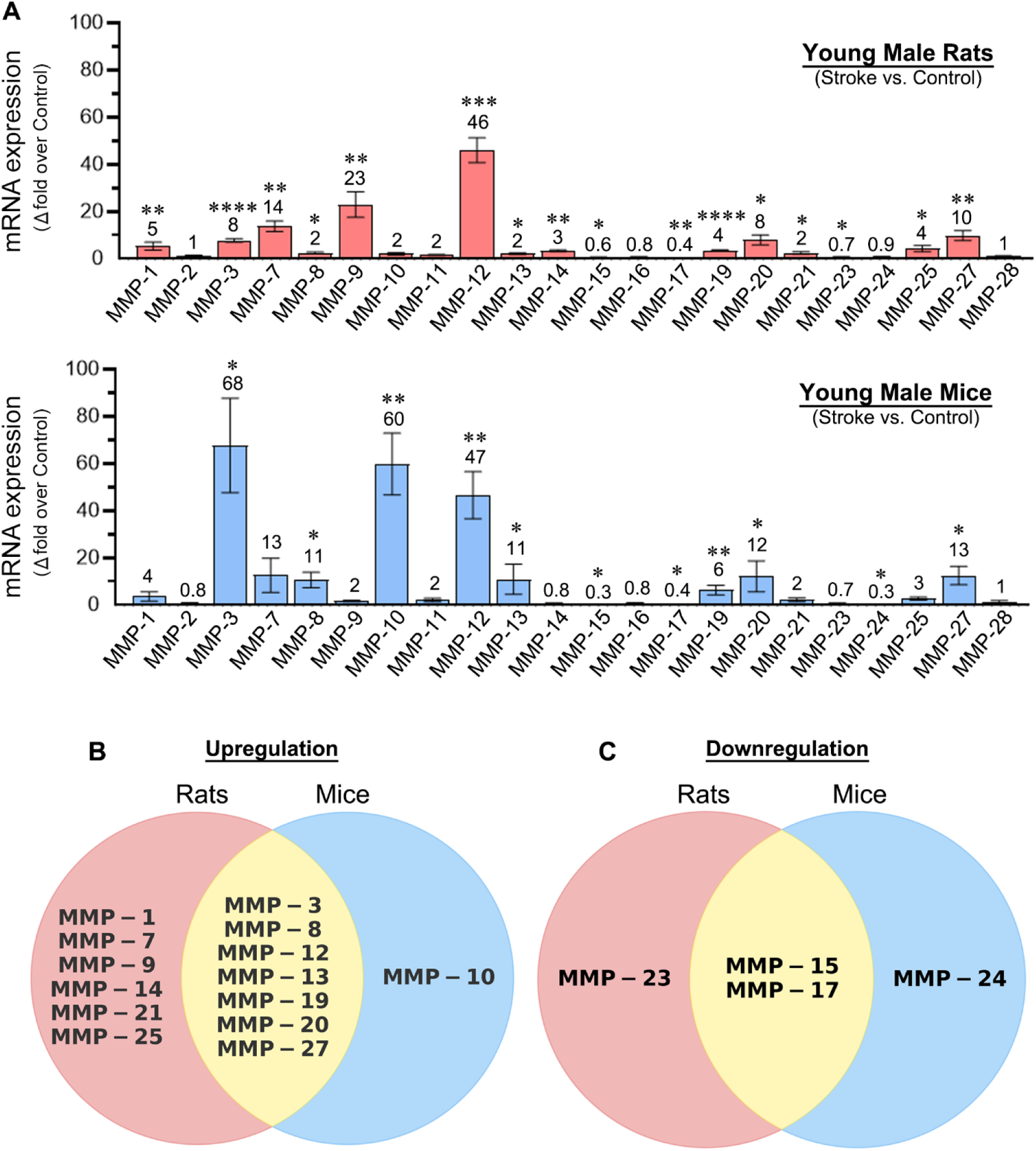
MMP expression in the ischemic brain of young male rats and mice. (**A**) Bar graphs show the quantified mRNA expression levels (fold change vs. the respective control group) for all MMPs in the ipsilateral (ischemic) hemisphere of young male rats and mice subjected to MCAO (2-h in rats and 1-h in mice) followed by reperfusion. Values above each bar indicate the fold change. Data are presented as mean ± SEM; n = 5–6/group. \**p* < 0.05, \*\**p* < 0.01, \*\*\**p* < 0.001, \*\*\*\**p* < 0.0001 vs. control group. (**B, C**) Venn diagrams illustrate significantly upregulated (**B**) and downregulated (**C**) MMPs (vs. controls) post-MCAO and reperfusion in the ipsilateral hemisphere of rats and mice, highlighting species differences across the 22 MMPs examined.

## 4. Discussion

The results of this study indicate that aging, sex, and species influence brain MMP expression patterns following cerebral I/R. These findings emphasize the importance of considering age, sex, and species when assessing and validating the effectiveness of novel treatments targeting selective MMPs, individually or in combination, in preclinical rodent stroke models. Overall MMP expression levels in the ischemic brain were higher in aged male mice and lower in aged female mice compared with sex-matched young mice. Although the expression of several MMPs in the ischemic brain changed with age and sex after cerebral I/R, MMP-3, MMP-10, and MMP-12 were the three most strongly upregulated MMPs in both male and female mice, regardless of age. By contrast, in young male rats, MMP-12 was the most prominently upregulated MMP in the ischemic brain followed by MMP-9 and MMP-7.

MMP-12 (metalloelastase) was the only MMP that was markedly upregulated in the ischemic brain following cerebral I/R across species and, within mice, across age groups and sexes. The substantial upregulation of MMP-12 observed in this study is consistent with our previous findings (Chelluboina and Warhekar et al., 2015; Nalamolu et al., 2018). Notably, baseline MMP-12 expression was significantly higher in young females, but not aged females, compared with age-matched males. In addition, aging increased baseline MMP-12 expression in males, but not females, compared with sex-matched young mice. Multiple resident brain cell types (e.g., microglia, oligodendrocytes, and neurons) as well as infiltrating leukocytes (including monocytes/macrophages and neutrophils) may contribute to MMP-12 production in the ischemic brain (Svedin et al., 2009; Chelluboina and Warhekar et al., 2015; Iyer et al., 2015; Hohjoh et al., 2020). Increased MMP-12 can degrade or process a broad range of extracellular matrix proteins and bioactive molecules, including type IV collagen, laminin, fibronectin, vitronectin, elastin, fibrillin-1, chondroitin sulfate and heparan sulfate proteoglycans, myelin basic protein, plasminogen, progranulin, α1-antitrypsin, tissue factor pathway inhibitor, N-cadherin, and pro-tumor necrosis factor-α, thereby contributing to brain damage and neurological deficits (Werb and Gordon, 1975; Shapiro et al., 1992; Shapiro et al., 1993; Chandler et al., 1996; Gronski et al., 1997; Dong et al., 1997; Cornelius et al., 1998; Ashworth et al., 1999; Belaaouaj et al., 2000; Chen, 2004; Dwivedi et al., 2009; Suh et al., 2012; Chelluboina and Warhekar et al., 2015). Consistent with this, reducing MMP-12 expression after cerebral I/R decreased BBB disruption, apoptosis, inflammation, and demyelination, and improved recovery of somatosensory, motor, and cognitive functions in young rodents (Chelluboina and Warhekar et al., 2015; Chelluboina and Klopfenstein et al., 2015; Arruri et al., 2022; Challa et al., 2022). In the present study, marked MMP-12 upregulation on post-ischemic day 1 across species and, within mice, across age groups and sexes, underscores the importance of initiating MMP-12-targeting therapies promptly after reperfusion. Given its detrimental effects and consistently strong induction, MMP-12 may represent a promising therapeutic target for acute ischemic stroke patients undergoing recanalization.

Following MMP-12, MMP-3 (stromelysin-1) and MMP-20 (enamelysin) were the only additional MMPs that were significantly upregulated after cerebral I/R across all groups examined. The clinical relevance of MMP-3 upregulation after ischemic stroke is supported by increased MMP-3 expression in infarcted brain tissue from human stroke patients (Cuadrado et al., 2009). Baseline MMP-3 expression was significantly higher in females than in age-matched males regardless of their age, whereas aging did not affect baseline MMP-3 expression in either sex. These findings indicate that both rats and mice are the appropriate preclinical models for investigating the role of MMP-3 in ischemic stroke. Potential cellular sources of MMP-3 in the ischemic brain include neurons, oligodendrocytes, and microglia/macrophages (Solé et al., 2004). Consistent with a detrimental role, MMP-3 knockout mice exhibit reduced infarct size and downregulated inflammation-associated gene signatures following cerebral I/R (Hamblin et al., 2024). Hyperglycemia further enhances MMP-3 activity after stroke, and its inhibition reduces edema, hemorrhagic transformation, and improves functional outcomes (Hafez et al., 2016; Abdul et al., 2023). Although MMP-3 appears to be a promising therapeutic target for ischemic stroke, the significance of elevated MMP-20 in the brain following cerebral I/R remains to be determined.

This study found that MMP-10 (stromelysin-2) was increased in the ischemic brain in both sexes of young and aged mice, but not in rats. The lack of a statistically significant change in MMP-10 expression in the rat ischemic brain is consistent with our earlier findings (Chelluboina and Warhekar et al., 2015). However, the robust upregulation of MMP-10 observed in the present study across young and aged mice of both sexes differs from our previous study, in which only young male mice were evaluated and MMP-10 was not increased but was instead decreased (Nalamolu et al., 2018). In contrast to our prior work, the present investigation used newly designed MMP-10 qPCR primers with an optimized annealing temperature, applied stringent exclusion criteria based on neurological deficit scores, and used a Doccol filament with a slightly larger coated tip diameter (6024 vs 6023) to induce MCAO. Collectively, these methodological refinements and the consistent MMP-10 upregulation across mouse groups suggest that the present findings are more robust and provide a more reliable estimate of post-stroke MMP-10 upregulation in mice. Notably, neither aging nor sex influenced baseline MMP-10 expression. MMP-10 upregulation after ischemic stroke is clinically relevant, as elevated MMP-10 expression has been reported in infarcted brain tissue from human stroke patients (Cuadrado et al., 2009). Consistent with our findings, mice may be a particularly informative species for evaluating MMP-10-targeted therapies, given that MMP-10 increased in mice but not rats in the present study. Based on literature, MMP-10 may play a dual role in ischemic stroke. MMP-10 knockout mice subjected to MCAO exhibited larger infarcts than wild-type mice, whereas administration of recombinant MMP-10 reduced infarct size in wild-type mice after MCAO (Orbe et al., 2011). In diabetic mice subjected to MCAO, administration of recombinant MMP-10 was reported to be more efficacious than tPA in reducing oxidative stress, BBB disruption, infarct size, and neurodegeneration, without increasing hemorrhage (Navarro-Oviedo et al., 2019). By contrast, serum pro-MMP-10 levels in patients with acute ischemic stroke have been associated with severe edema, larger infarct volume, worse neurological deficits, and poor functional outcome (Rodríguez et al., 2013). Therefore, additional studies are needed to define the net effects of elevated endogenous MMP-10 in the ischemic brain of young and aged rodents of both sexes following cerebral I/R on a range of post-stroke outcomes.

MMP-9 (gelatinase B) was the second most strongly upregulated MMP in rats after cerebral I/R. The significant upregulation of MMP-9 in the rat ischemic brain observed in the present study is consistent with our earlier findings (Chelluboina and Warhekar et al., 2015). Post-stroke elevation of MMP-9 is clinically relevant, as increased MMP-9 expression has been reported in infarcted brain tissue from human stroke patients (Cuadrado et al., 2009). Interestingly, despite being one of the most widely studied MMPs in ischemic stroke, MMP-9 was not upregulated in mice, except in aged males, where the increase reached statistical significance. In contrast, our prior study reported an approximately 24-fold upregulation of MMP-9 in the ischemic brain of young male mice on post-ischemic day 1 (Nalamolu et al., 2018). Despite this discrepancy in males, neither young nor aged females exhibited elevated MMP-9 expression in the ischemic brain. It is possible that MMP-9 upregulation in mice occurs primarily during the early hours after reperfusion. Taken together, these observations suggest that rats may represent a more sensitive and clinically relevant rodent species for evaluating MMP-9-targeted therapies in ischemic stroke. Aging did not affect baseline MMP-9 expression in either sex. However, baseline MMP-9 expression was significantly higher in aged females than in age-matched males, whereas no sex difference was observed in young mice. In contrast to the potential dual role of described for MMP-10, MMP-9 exhibits a predominantly detrimental role in ischemic stroke. Elevated serum MMP-9 levels during the acute phase of ischemic stroke have been associated with increased mortality and severe disability (Zhong et al., 2017). Numerous studies have demonstrated increased MMP-9 expression in the ischemic brain of rats and mice and shown that its inhibition or genetic deletion reduces edema, neuroinflammation, BBB disruption, and infarct volume while improving functional recovery (Asahi et al., 2001; Sumii and Lo, 2002; Aoki et al., 2002; Montaner et al., 2003; Horstmann et al., 2003; Lo et al., 2004; Ning et al., 2006; Sandoval and Witt, 2008; Hu et al., 2009; Hu et al., 2011; Du et al., 2025). Collectively, these findings support MMP-9 as a promising therapeutic target in acute ischemic stroke, particularly in the context of reperfusion, given its detrimental role and its upregulation in the ischemic brain of humans and rats.

Overall, MMP expression increased substantially in the ischemic brain during the acute phase following cerebral I/R, regardless of age, sex, or species. Broad-spectrum (non-selective) MMP inhibition has been shown to reduce BBB disruption and infarct volume and to improve functional outcomes in experimental models of ischemic stroke (Kawa et al., 2025). However, although non-selective MMP inhibition may yield net beneficial effects, it may also suppress MMPs with potentially beneficial roles, such as MMP-10. An ideal therapeutic strategy would selectively inhibit detrimental MMPs while sparing those that contribute to repair and recovery. Moreover, the optimal duration of MMP inhibition remains uncertain, as prolonged suppression may interfere with recovery processes mediated by certain MMPs.

This study has several limitations. First, we assessed only mRNA expression of MMPs and did not measure their protein levels. Therefore, the extent to which mRNA changes translate to changes in protein expression remains to be determined. Second, to investigate similarities and differences between species, we profiled MMP expression only in the ischemic brain of young male rats. We anticipate that the overall MMP expression profile in young female rats would be broadly comparable to that in young male rats as in mice, although the relative magnitude of induction may differ across MMPs. Aged rats were not included in this study because they are less frequently used than mice in preclinical studies evaluating the efficacy of novel stroke therapies. Third, MMP expression was assessed only on post-ischemic day 1. Because MMP expression can remain elevated during both acute and chronic phases after cerebral I/R (Chelluboina and Warhekar et al., 2015), MMPs induced early may be more closely linked to acute brain injury, whereas those induced later may contribute to repair and recovery. Future research should examine sex-, age-, and species-related differences in MMP expression at additional time points, including at least one during the recovery phase, and should also evaluate corresponding MMP protein expression.

In summary, the findings of this study indicate that brain MMP expression following cerebral I/R is modulated by age, sex, and species, underscoring the importance of considering these factors when targeting MMPs, individually or in combination, in preclinical rodent stroke models. Notably, among all MMPs examined, MMP-12 showed the most prominent upregulation across species and, within mice, across age groups and sexes, suggesting that MMP-12 may represent a promising therapeutic target for acute ischemic stroke in the context of reperfusion. However, the extent to which MMP-12 is upregulated in human brain tissue after acute ischemic stroke, particularly in patients who undergo recanalization, remains largely uncharacterized. Future studies defining MMP expression patterns in human ischemic brain tissue would provide critical translational context and help guide the selection of the most appropriate rodent species for evaluating MMP-targeted therapies, thereby improving clinical translatability and potentially minimizing failures in clinical trials.

## Author contributions

Conceptualization, funding acquisition, and supervision: K.K.V. and J.D.K.; Animal surgeries and care: S.R.C., K.K.V., and I.M.B.; Methodology, investigation, and data acquisition: S.R.C., V.V., S.N.J., I.M.B., N.K., P.U., and S.R.M.; Data analysis and interpretation: K.K.V., C.A.F., and S.R.C.; Manuscript preparation: K.K.V.; Manuscript editing: C.A.F.; Manuscript review and final approval: All authors.

## Acknowledgments

We thank the National Institutes of Health (NIH) for financial support.

## Data availability

All data supporting the key findings of this study are included in this article. Additional raw data of real-time PCR, including instrument-generated data files and associated analyses, are available from the corresponding author upon reasonable request.

