## Supplementary Fig. 1 for "Impact of Aging, Sex, and Species on the mRNA Expression of Matrix Metalloproteinases Following Ischemic Stroke"

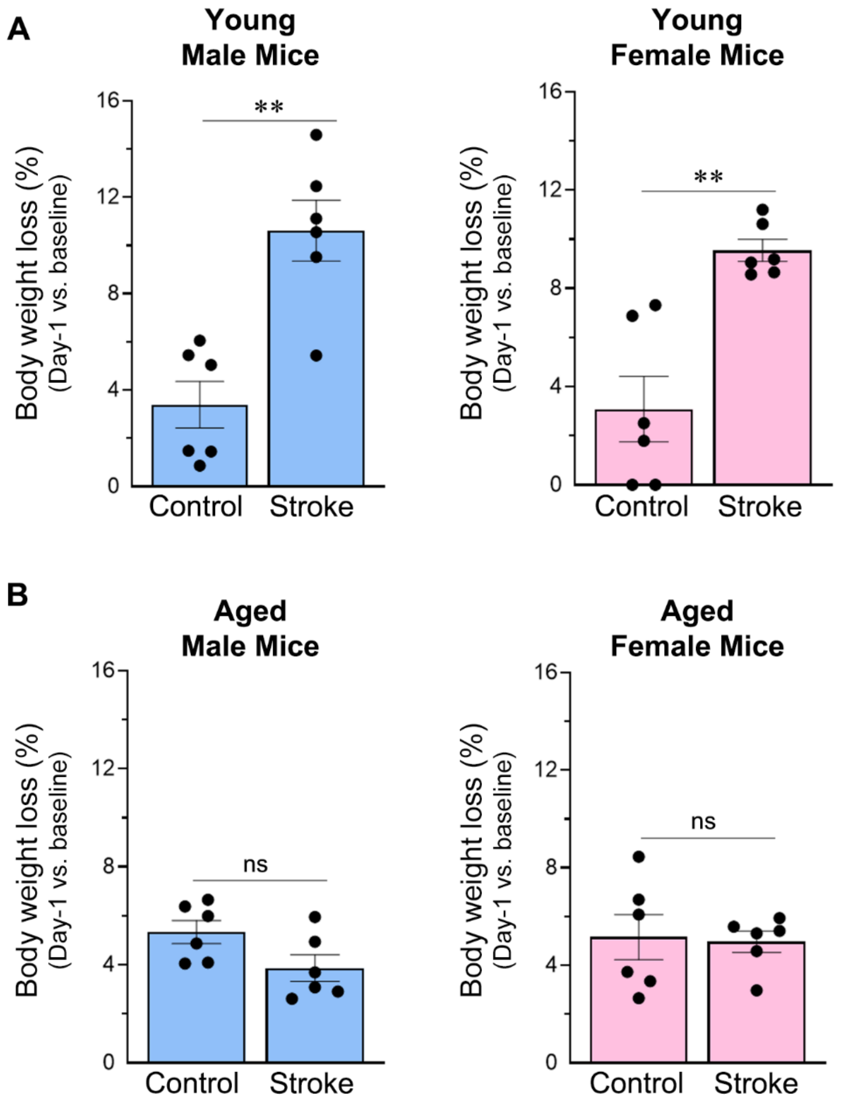


**Supplementary Fig. 1. Post-stroke body weight loss in young and aged mice of both sexes.** Column scatter plots show the percent body weight loss on post-ischemic day 1 after 1-h MCAO (relative to baseline body weight) in (**A**) young and (**B**) aged male and female mice. $^{**}p < 0.01$; ns, not significant.
