## Supplementary Fig. 2 for "Impact of Aging, Sex, and Species on the mRNA Expression of Matrix Metalloproteinases Following Ischemic Stroke"

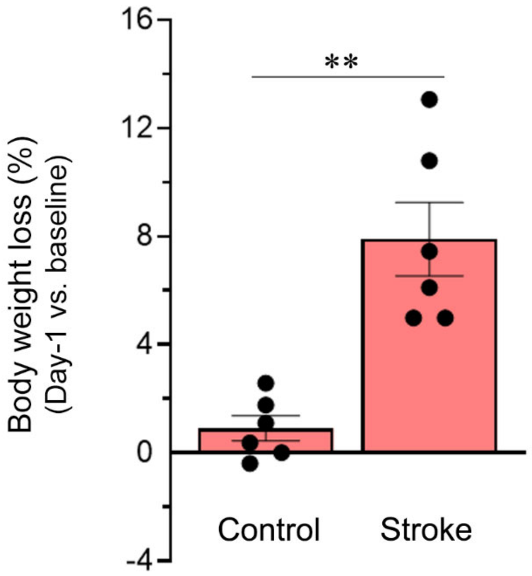


**Supplementary Fig. 2. Post-stroke body weight loss in young male rats.** Column scatter plot shows the percent body weight loss on post-ischemic day 1 after 2-h MCAO (relative to baseline body weight) in young male rats. **$p < 0.01$.
