## Supplementary Fig. 3 for "Impact of Aging, Sex, and Species on the mRNA Expression of Matrix Metalloproteinases Following Ischemic Stroke"

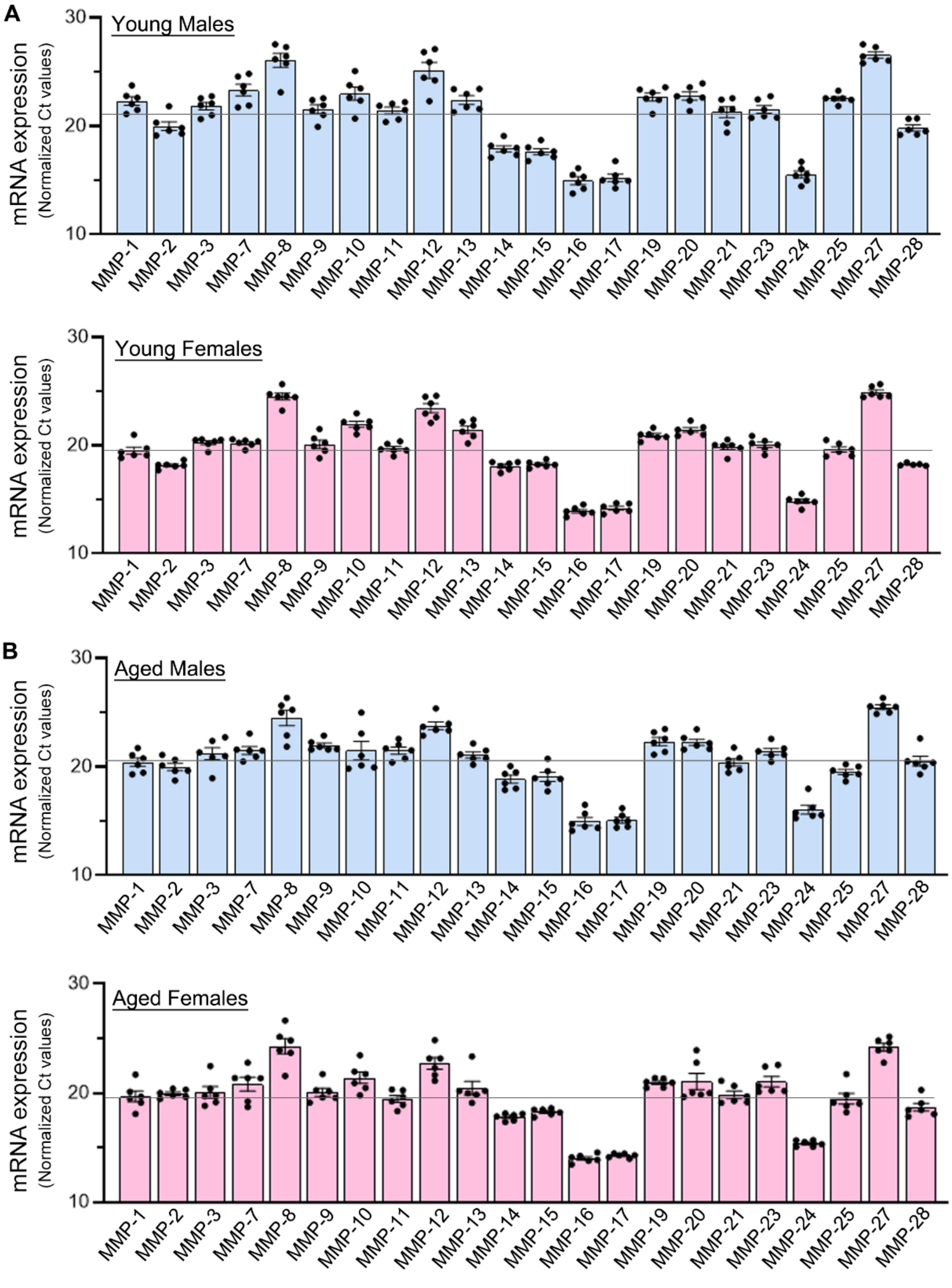


**Supplementary Fig. 3. Normalized mRNA expression (Ct values) of MMPs in the ischemic brain of young and aged mice of both sexes.** Column scatter plots show normalized Ct values (normalized to the internal standard, *18S* rRNA) for all MMPs in the ipsilateral (ischemic) hemisphere of (**A**) young and (**B**) aged male and female mice (n=5-6/group). The horizontal line in each group indicates the mean normalized Ct value across the 22 MMPs examined.
