## Supplementary Table 1 for "Impact of Aging, Sex, and Species on the mRNA Expression of Matrix Metalloproteinases Following Ischemic Stroke"

**Supplementary Table 1.** Experimental groups, description, and animal numbers.

| Group # | Group description | Number of animals | | | | |
|---|---|---|---|---|---|---|
| | | Included | Excluded | | | Total |
| | | | Mortality | Hemorrhage | mNSS<8 NDS<2 | |
| *Control groups* | | | | | | |
| R1 | Young male rats subjected to MCAO surgery (no suture inserted). | 6 | 0 | 0 | 0 | 6 |
| M1 | Young male mice subjected to MCAO surgery (no suture inserted). | 6 | 0 | 0 | 0 | 6 |
| M3 | Young female mice subjected to MCAO surgery (no suture inserted). | 6 | 0 | 0 | 0 | 6 |
| M5 | Aged male mice subjected to MCAO surgery (no suture inserted). | 6 | 0 | 0 | 0 | 6 |
| M7 | Aged female mice subjected to MCAO surgery (no suture inserted). | 6 | 0 | 0 | 0 | 6 |
| *Stroke groups* | | | | | | |
| R2 | Young male rats subjected to 2-h MCAO | 6 | 0 | 0 | 2 | 8 |
| M2 | Young male mice subjected to 1-h MCAO | 6 | 2 | 2 | 0 | 10 |
| M4 | Young female mice subjected to 1-h MCAO | 6 | 0 | 0 | 1 | 7 |
| M6 | Aged male mice subjected to 1-h MCAO | 6 | 4 | 1 | 0 | 11 |
| M8 | Aged female mice subjected to 1-h MCAO | 6 | 1 | 1 | 0 | 8 |

mNSS, modified neurological severity score; NDS, neurological deficit score; MCAO, middle cerebral artery occlusion
