## Supplementary Table 2 for "Impact of Aging, Sex, and Species on the mRNA Expression of Matrix Metalloproteinases Following Ischemic Stroke"

**Supplementary Table 2.** Rat primers used for real time PCR analysis.

| Gene | NCBI Reference Sequence | Primer Sequence | |
|---|---|---|---|
| | | **Forward (5' - 3')** | **Reverse (5' - 3')** |
| MMP-1 | NM_001134530 | cagcagttatttgggctgaaag | tttggtccaacgaggattgt |
| MMP-2 | NM_031054 | gctgatactgacactggtactg | tgtcactgtccgccaaataa |
| MMP-3 | NM_133523 | ggaccagggattaatggagatg | agcattggctgagtgaaaga |
| MMP-7 | NM_012864 | gagtgccagatgttgcagaa | ctgcagtcccccaactaa |
| MMP-8 | NM_022221 | tgggctctaagtgcctatga | cgtatctccagcattggttgt |
| MMP-9 | NM_031055 | cactgtaactgggggcaact | cacttcttgtcagcgtcgaa |
| MMP-10 | NM_133514 | tcccaccgtgaagaagattg | cagtctcgggaagcctttat |
| MMP-11 | NM_012980 | gctgagggctatgcctact | gaagcccaggtccacaaata |
| MMP-12 | NM_053963 | gctagaagtaactgggcaact | gagataccgcttcatccatctt |
| MMP-13 | NM_133530 | gccctgatgtttcccatctat | ggtcacacttctctggtgttt |
| MMP-14 | NM_031056 | gtaccccaagtcagctct | cagtgaacgctggcagtaaa |
| MMP-15 | NM_001106168 | aacgtcaggatgggcattt | tgtgtcaatgcggtcatagg |
| MMP-16 | NM_080776 | cagctctggaagaaggtt | gcctgactgcatggtctct |
| MMP-17 | NM_001105925 | ctgtactggcgctatgacga | ctgcctctagctcaccat |
| MMP-19 | NM_001107159 | acttgggcattcccgatatac | acctcttcctcctcatcttct |
| MMP-20 | NM_001106800 | ggtcctcctcgcacaattta | agctgtagtattcctctccca |
| MMP-21 | NM_001106308 | gggtcatgtgagggatcattt | agctgttgcggaagaagtag |
| MMP-23 | NM_053606 | ctgatgcactcacagcaaga | gcccaggatgtacacacaaa |
| MMP-24 | NM_031757 | tgagcaggaggaggagaaata | tcgaaagcctgacgaatagc |
| MMP-25 | NM_001427286 | tggagtatccagtccttctct | tgagaatccacctcttggaac |
| MMP-27 | NM_001106799 | caggcatatctcaaccagttct | tttcccgtcactgtcaatcc |
| MMP-28 | NM_001079888 | gcaaagccaggtcacaaatg | gcgctgacacattactccata |
| *18S* rRNA | NR_046237 | acgtctgccctatcaactttc | ttggatgtggtagccgtttc |

PCR, polymerase chain reaction; MMP, matrix metalloproteinase
