## Supplementary Table 3 for "Impact of Aging, Sex, and Species on the mRNA Expression of Matrix Metalloproteinases Following Ischemic Stroke"

**Supplementary Table 3.** Mouse primers used for real time PCR analysis.

| Gene | NCBI Reference Sequence | Primer Sequence Forward (5' - 3') | Primer Sequence Reverse (5' - 3') |
|---|---|---|---|
| MMP-1 | NM_032006 | ttgggctcactcattctagtg | tgttggctggatgggattt |
| MMP-2 | NM_008610 | gtcgcccctaaaacagacaa | ggtctcgatggtgttctggt |
| MMP-3 | NM_010809 | cagacttgtcccgtttccat | ggtgctgactgcatcaaaga |
| MMP-7 | NM_010810 | tcggatcgtagtggatcaaatc | tctccttgcgaagccaatta |
| MMP-8 | NM_008611 | gccttcccagtacctgaaca | actccacatcgaggcatttc |
| MMP-9 | NM_013599 | cgtcgtgatccccacttact | aacacacagggtttgccttc |
| MMP-10 | NM_019471 | aagctggactccaacactatg | cctgtaggtgatgtgggattt |
| MMP-11 | NM_008606 | gcatgcagctctgcctaata | gttccgggcattcagtacat |
| MMP-12 | NM_001320076 | ccaagcatcccatctgctat | ggtcaaagacagctgcatca |
| MMP-13 | NM_008607 | gttgacaggctccgagaaat | catcaggcactccacatctt |
| MMP-14 | NM_008608 | ccgccatgcaaaagttctat | gcccaccttaggggtgtaat |
| MMP-15 | NM_008609 | cccacaggtcacaccttctt | ccagtacttggtgcccttgt |
| MMP-16 | NM_019724 | ggagacagttccccatttga | ccagctcatggactgctaca |
| MMP-17 | NM_011846 | ggagcaagaggaacctttctt | ggaagttcaagggtgtgatgt |
| MMP-19 | NM_021412 | agacaagagatgaggaggaaga | agtcgcccttgaaagcataa |
| MMP-20 | NM_013903 | ggtcctccacggacaatttat | agctgtagtattcctctccca |
| MMP-21 | NM_152944 | agagatacctagcccaaagga | acagcgtggcttgttcata |
| MMP-23 | NM_011985 | gaccacttcaacctcacataca | ccacctcacggaaactgaat |
| MMP-24 | NM_010808 | tgagcaggaggaggagaaata | tcgaaagcctgacgaatagc |
| MMP-25 | NM_001033339 | ttgagcccgcagattgtt | cccgagcaaagtggataatga |
| MMP-27 | NM_001310717 | agatggcacaagcaatgga | cctcgacccatggaagaaatag |
| MMP-28 | NM_001320300 | tacaagcagcatctttcctacc | actccagtgctgacacattac |
| *18S* rRNA | NR_003278 | tgagaaacggctaccacatc | gcctcgaaagagtcctgtatt |

PCR, polymerase chain reaction; MMP, matrix metalloproteinase
