## Supplementary Table 4 for "Impact of Aging, Sex, and Species on the mRNA Expression of Matrix Metalloproteinases Following Ischemic Stroke"

**Supplementary Table 4.** Modified Neurological Severity Scores in rats subjected to 2-h MCAO.

| Group # | Age/Sex | Animal ID | Modified Neurological Severity Score (mNSS) |
|---|---|---|---|
| R2 | Young male rats | SDRM - 305 | 8 |
| | | SDRM - 307 | 8 |
| | | SDRM - 308 | 8 |
| | | SDRM - 317 | 10 |
| | | SDRM - 324 | 12 |
| | | SDRM - 328 | 12 |

MCAO, middle cerebral artery occlusion.
