## Supplementary Table 5 for "Impact of Aging, Sex, and Species on the mRNA Expression of Matrix Metalloproteinases Following Ischemic Stroke"

**Supplementary Table 5.** Neurological Deficit Scores in mice subjected to 1-h MCAO.

| Group # | Age/Sex | Animal ID | Neurological Deficit Score (NDS) |
|---|---|---|---|
| M2 | Young male mice | C57MY - 50 | 2 |
| | | C57MY - 56 | 3 |
| | | C57MY - 58 | 2 |
| | | C57MY - 59 | 3 |
| | | C57MY - 64 | 2 |
| | | C57MY - 65A | 3 |
| M4 | Young female mice | C57FY - 11 | 3 |
| | | C57FY - 30 | 3 |
| | | C57FY - 35 | 3 |
| | | C57FY - 36 | 3 |
| | | C57FY - 37 | 3 |
| | | C57FY - 38 | 3 |
| M6 | Aged male mice | C57MA - 74 | 3 |
| | | C57MA - 83 | 3 |
| | | C57MA - 86 | 3 |
| | | C57MA - 88 | 3 |
| | | C57MA - 92 | 3 |
| | | C57MA - 95 | 3 |
| M8 | Aged female mice | C57FA - 56 | 3 |
| | | C57FA - 57 | 2 |
| | | C57FA - 62 | 3 |
| | | C57FA - 63 | 3 |
| | | C57FA - 64 | 2 |
| | | C57FA - 65 | 3 |

MCAO, middle cerebral artery occlusion.
